# Plastid phylogenomics resolves ambiguous relationships within the orchid family and provides a solid timeframe for biogeography and macroevolution

**DOI:** 10.1101/774018

**Authors:** Maria Alejandra Serna-Sánchez, Oscar A. Pérez-Escobar, Diego Bogarín, María Fernanda Torres, Astrid Catalina Alvarez-Yela, Juliana E. Arcila, Climbie F. Hall, Fábio de Barros, Fábio Pinheiro, Steven Dodsworth, Mark W. Chase, Alexandre Antonelli, Tatiana Arias

## Abstract

Recent phylogenomic analyses based on the maternally inherited plastid organelle have enlightened evolutionary relationships between the subfamilies of Orchidaceae and most of the tribes. However, uncertainty remains within several subtribes and genera for which phylogenetic relationships have not ever been tested in a phylogenomic context. To address these knowledge-gaps, we here provide the most extensively sampled analysis of the orchid family to date, based on 78 plastid coding genes representing 264 species, 117 genera, 18 tribes and 28 subtribes. Divergence times are also provided as inferred from strict and relaxed molecular clocks and birth-death tree models. Our taxon sampling includes 51 newly sequenced plastid genomes produced by a genome skimming approach. We focus our sampling efforts on previously unplaced clades within tribes Cymbidieae and Epidendreae. Our results confirmed phylogenetic relationships in Orchidaceae as recovered in previous studies, most of which were recovered with maximum support (209 of the 262 tree nodes). We provide for the first time a clear phylogenetic placement for Codonorchideae within subfamily Orchidoideae, and Podochilieae and Collabieae within subfamily Epidendroideae. We also identify relationships that have been persistently problematic across multiple studies, regardless of the different details of sampling and genomic datasets used for phylogenetic reconstructions. Our study provides an expanded, robust temporal phylogenomic framework of the Orchidaceae that paves the way for biogeographical and macroevolutionary studies.

## 1. Introduction

Orchidaceae, with *ca*. 25,000 species and ∼800 genera^1,2^ are one of two of the most diverse and widely distributed flowering plant families on Earth and have captivated the attention of scientists for centuries^3^. The family has a striking morphological and ecological diversity and evolved complicated interactions with fungi, animal and other plants^4,5^ and a diverse array of sexual systems^6–8^. Numerous efforts have been made to understand the natural history, evolution and phylogenetic relationships within the family^2,7,9–13^. To date, there are seven nuclear genome sequences available, i.e., *Apostasia shenzhenica*^14^, *Dendrobium catenatum*^15^, *D. officinale*^16^, *Gastrodia elata*^17^, *Phalaenopsis equestris*^18^, a *Phalaenopsis* hybrid cultivar^19^, *P. aphrodite*^20^, *Vanilla planifolia*^21^, 221 complete plastid genomes and 2,678 sequence read archives for Orchidaceae in NCBI (accessed 22 August 2020).

Phylogenomic approaches have been implemented to infer relationships between major orchids clades in deep and recent time^2,10,12,13,22,23^, but extensive uncertainties remain regarding the phylogenetic placement of several subtribes. This knowledge-gap stems from the large gaps in both taxonomic and genomic sampling efforts that would be required to comprehensively cover all major orchid clades (subtribes/groups of genera). Givnish et al.^2^ published the first well-supported analysis of Orchidaceae based on plastid phylogenomics. They performed a maximum likelihood (ML) analysis of 75 genes from the plastid genome of 39 orchid species, covering 22 subtribes, 18 tribes and five subfamilies. This robust but taxonomically under-sampled study agreed corroborated relationships of the subfamilies and tribes, observed in previous studies^10–13^.

Previous orchid studies have failed to resolve relationships in rapidly diversifying clades^24–26^ because of reduced taxon and data sampling^27^. This is particularly true for Cymbidieae and Pleurothallidinae, the two most species-rich groups in which generic relationships are largely the product of rapid diversification^28^ that is difficult to resolve using only a few loci^25,29^. Cymbidieae comprise 10 subtribes, ∼145 genera and nearly 3,800 species^1^, 90% of which occur in the Neotropics ^28^. Four of these subtribes are among the most species-rich in the Andean and Chocoan region (Maxillariinae, Oncidiinae, Stanhopeinae and Zygopetaliinae^30,31^). Pleurothallidinae include ∼ 5,500 exclusively Neotropical species in 47 genera. Pleurothallid orchids are one of the most prominent components of the cloud forest flora in the northern and central Andes and Central America^32^.

Another group in which phylogenetic relationships are unresolved is Orchidoideae ^1,33^. This group comprises four mostly terrestrial tribes, 25 subtribes and over 3,600 species. The subfamily occurs on all continents except the Antarctic. Previous efforts to disentangle the phylogenetic relationships in the subfamily have mostly relied on a small set of nuclear and plastid markers^34^, and more recently on extensive plastid coding sequence data^2^.

The wide geographical range of these groups in the tropics and temperate regions and their striking vegetative and reproductive morphological variability make them ideal model clades for disentangling the contribution of abiotic and biotic drivers of orchid diversification across biomes. Occurring from alpine ecosystems to grasslands, they have conquered virtually all ecosystems available in any elevational gradient^35–37^, showing independent transitions to terrestrial, rupicolous and epiphytic habit. Moreover, they have evolved a diverse array of pollination systems^38–40^, rewarding species offering scent, oil and nectar, and even food- and sexual deceptive species^41,42^. However, the absence of a solid phylogenetic framework has precluded the study of how such systems evolved and the diversification dynamics of Cymbidieae, Pleurothallidinae and Orchidoideae more broadly.

Phylogenetic analyses are crucial to understanding the drivers of diversification in orchids, including the mode and tempo of morphological evolution^30,43^. High-throughput sequencing and modern comparative methods have enabled the production of massive molecular datasets to reconstruct evolutionary histories and thus provide unrivalled knowledge on plant phylogenetics^44^. Here, we present the most densely sampled plastid analysis of Orchidaceae, including data from 51 newly sequenced plastid genomes,. We apply two general approaches: a) maximum likelihood phylogenetic analysis conducted on 78 plastid coding regions to inform relationships; b) Bayesian inference in combination with strict and relaxed molecular clocks and a birth-death model applied to a subset of the plastid coding regions to produce a temporal framework of the orchid family. Our study expands the current generic representation for the Orchidaceae and clarifies previously unresolved phylogenetic relations within the Cymbidieae, Pleurothallidinae and Orchidoideae. The results reported here provide a robust framework for the orchid family and new insights into relationships at both deep and shallow phylogenetic levels.

## 2. Results

### 2.1 Phylogenetic relationships and divergence times in the orchid family

The ML tree derived from the 78 plastid genes is provided in Fig. 1. Two hundred-and-thirty-one nodes were recovered as strongly supported (i.e. likelihood bootstrap percentage [LBP] = 85-100), of which 209 attained maximum support. Only 26 nodes recovered LBPs between 25 and 84 (Fig. 1, inset). Unsupported relationships were restricted to Epidendroideae and Orchidoideae but were more frequent in Epidendroideae and often linked to low levels of sequence variation. Here, poorly supported relationships occurred mostly towards the backbone of the tribes Arethuseae, Cymbidieae, Epidendreae and Neottieae and Tropidieae + Nervilieae and the most recent common ancestor (MRCA) of Arethuseae, Malaxideae, Podochilieae, Collabieae, Epidendreae, Vandeae and Cymbidieae. Intrageneric relationships were robustly supported, with only two instances for which few nodes were recovered as poorly supported (*Dendrobium*: 3; *Cymbidium:* 1; Fig. S1).

**Figure 1.**
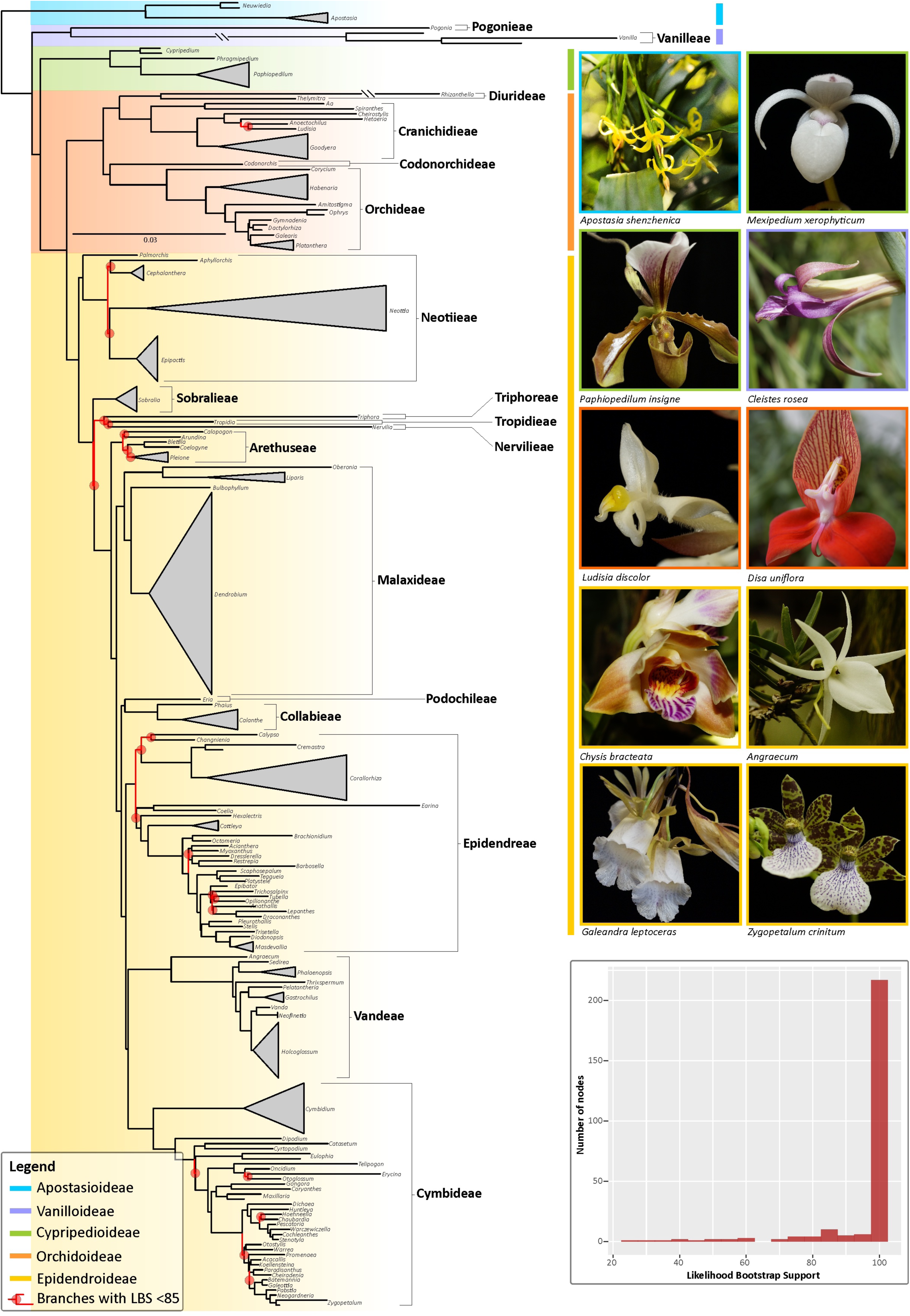
Maximum Likelihood phylogeny of the orchid family inferred from 78 coding plastid genes. Likelihood bootstrap support values (LBS) < 85% at nodes are highlighted in red together with their corresponding subtending branches. Orchid genera, tribes and subfamilies are indicated in the phylogeny together with photographs of selected representative species per subfamily. (Inset): Bar plot showing the frequency of LBS values at nodes as computed by bin intervals of 5 units.

Absolute times of divergence under strict and relaxed clocks for Orchidaceae, subfamilies and most tribes are provided in Table 1 (phylogenetic trees with mean ages and intervals of confidence produced under both clock models are provided on Figs S2, S3). Strict and relaxed molecular clocks revealed similar ages of divergence for the majority of the MRCAs of main orchid clades, although we found stark differences in the length of the 95% highest posterior density intervals (HPD) derived from both models are obvious, with the relaxed clock producing larger HPDs (Tab. 1; Fig. S2, S3; Tab. S1, S2). Under the strict and relaxed clocks, Orchidaceae diversified first during the late Cretaceous (88.1 my ± 3; 89.1 my ±9, respectively). The largest differences on the MRCA ages occurred in Epidendroideae (44 my ± 2 vs 60 my ± 10 under a strict and relaxed clock models, respectively) and Vanilloideae (80 my ± 4 vs 67 my ± 9). A complete account of mean and median ages, HPDs, branch lengths and rate values estimated for all nodes of chronograms estimated strict and relaxed molecular clock models are provided on Tab. S1, S2.

### 2.2. Phylogenetic informativeness of plastid genes

Phylogenetic informativeness plots are provided on Fig. S4 (see Tab. S3,S4 for a detailed account of PI per-site and net values for each assessed locus). Per-site and net phylogenetic informativeness (PI) analyses recovered both *ycf*1 as the most informative locus, which attained the highest values at a reference time (phylogenetic depth) of 0.51. On average, plastid loci attained their highest PI value at a reference time of 0.85 (SD=0.16). In contrast, the highest PI values of the 10 most informative loci occurred at an average reference time of 0.63 (SD=0.11) and 0.80 (SD=0.17) for per-site and net PI calculations.

## 3. Discussion

### 3.1 A robust temporal phylogenomic framework for the orchid family

Previous phylogenomic studies of the orchid family included up to 74 species representing 18 tribes, 19 subtribes and 66 genera^27^. Our study sampled 264 species from all subfamilies, representing 18 tribes (out of 22), 28 subtribes (out of 46) and 74 genera (∼10% of the currently recognised genera; Fig. 2). In general, our phylogenomic frameworks are in agreement with previously published family-wide orchid analyses either inferred from dozens of markers^2,13^ or from a handful of loci^29^. Here, representativeness of Cymbidieae and Epidendreae, two of the most prominent tropical Epidendroideae^45^clades, have increased from eight to 32 genera and six to 30, respectively^2,27^. In particular, relationships inferred from extensive plastid data within Zygopetaliinae (Cymbidieae) and Pleurothallidinae (Epidendreae) are presented for the first time. Our 78-coding sequence plastid ML analysis led to similar results as reported by Givnish et al.^2^, Niu et al.^13^ and Li et al.^27^ but with an overall clear increase in support: 22% of nodes with LBS < 85 in Givnish et al.^2^ and 21% in Li et al.^27^ *vs* 11.5% in this study. This is particularly evident in relationships inferred within Orchidoideae (see section 3.5 of *Discussion*) and Cymbidieae, Epidendreae (see sections 3.3 and 3.4 of *Discussion*, respectively) and Collabieae. For the last, for the first time we provide high support for the previously unresolved relationship of Podochilieae+Collabieae^2,27^.

**Figure 2.**
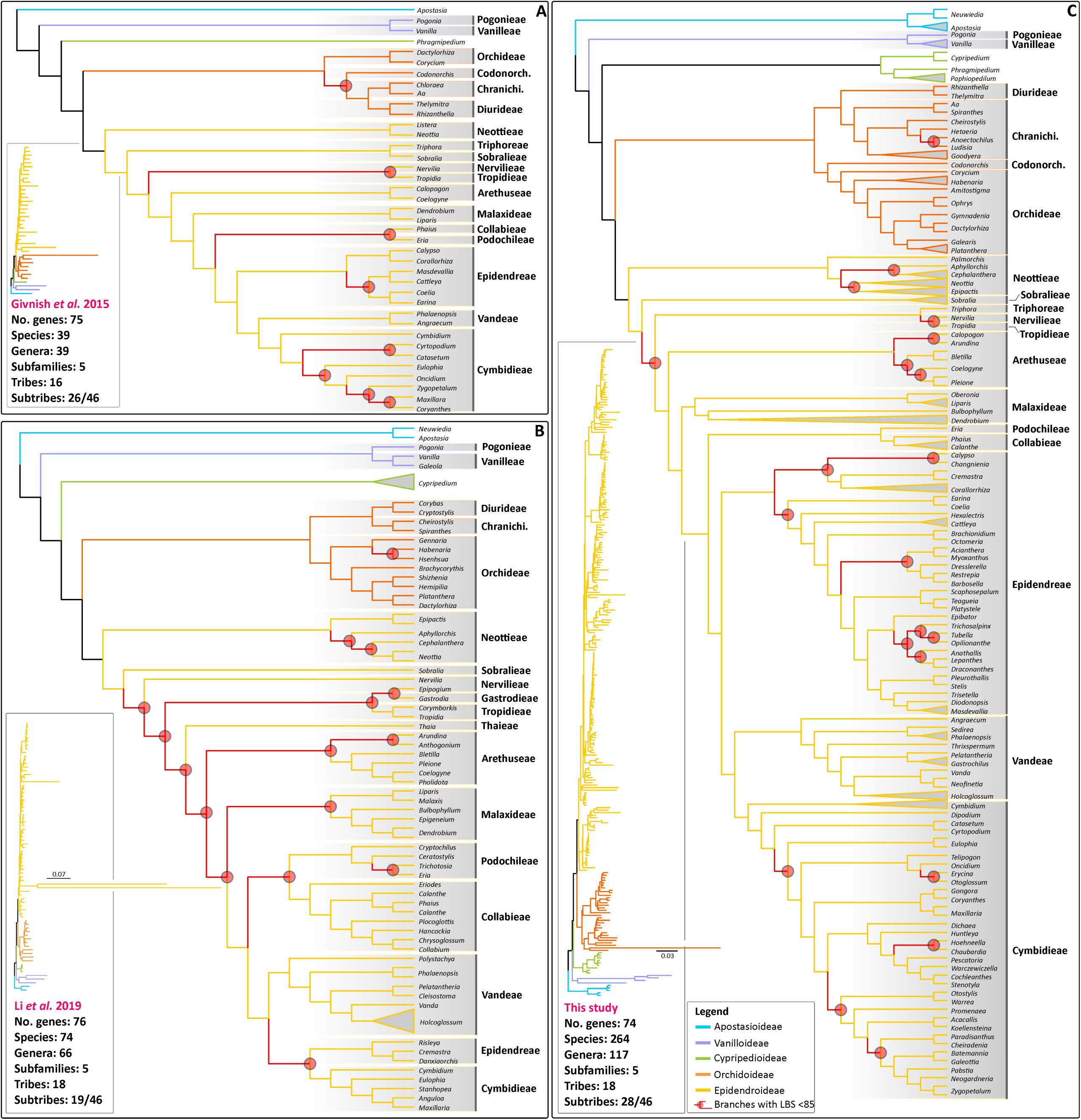
A comparison of the main plastid topologies of the orchid family published to date. A) Givnish et al.^2^’ inference based on 75 plastid genes and 39 orchid species; B) Li et al.^27^’ inference based on 76 plastid genes and 76 orchid species; C) This study: 78 plastid and 264 orchid species. LBP at nodes are highlighted in red together with their corresponding subtending branches. (Inset): trees witih branch lengths proportional to substitutions/site. Photos: O. Pérez-Escobar.

The absolute age estimates derived from our strict and relaxed molecular clocks and five of the most informative plastid loci are in line with previous nuclear-plastid multi-locus and phylogenomic plastid-only chronograms^2,46,47^. Nonetheless, our ML tree also identifies intricate relationships that have been consistently recovered as unsupported in several studies. These include unsupported basal nodes in Epidendroideae representing Sobralieae, Nervilieae and Tiphoreae^27,48^, *Arundina*+remainder of Arethuseae^27^, and the position of Eulophiinae in the Cymbidieae^25,28,49^ (Fig. 2). Uncertainty around the phylogenetic position of these clades might be due to limited taxon sampling in this and previous studies. Alternatively, intragenomic conflict^50–52^ and lack of phylogenetic informativeness required to sort out relationships derived from rapid diversifications^22,53,54^ in plastid DNA sequences (regardless of whether whole plastid genome datasets are employed^55^) might hamper the phylogenetic placement of clades with robust support.

### 3.3 Improved support of phylogenetic relationships within Cymbidieae

Multiple studies have inferred evolutionary relationships in Cymbidieae from morphological and molecular characters^28,29^. Relationships among subtribes have recently been estimated using the plastid genes *psaB, rbcL, matK* and *ycf1* combined with the low-copy nuclear gene *Xdh*^25^. Here, Cymbidiinae was sister to the remainder of Cymbidieae. Poorly supported and incongruent relationships were found among Catasetinae, Eulophiinae and Eriopsidinae, however, when compared with the topologies obtained by Whitten et al.^29^, Freudenstein & Chase^48^ and Pérez-Escobar et al.^7^

The most complete taxonomic sampling conducted to date under a plastid phylogenomic framework^2^ included 8 of 11 subtribes of Cymbidieae, but some inter-subtribal relationships were unresolved: Stanhopeinae (20 genera), Maxillariinae (12 genera), Zygopetalinae (36 genera), Oncidiinae (65 genera) and Eulophiinae (13 genera). A clade formed by Stanhopeinae and Maxillariinae had poor support (LBP=62) and their relationship to Zygopetaliinae also had low support (LBP=72). The relationship between Eulophiinae and a clade of Stanhopeinae, Maxillariinae, Zygopetalinae and Oncidiinae also had poor support (LBP=42). One of the outcomes of our expanded sampling (nine subtribes) is the improvement of support in Cymbidieae, more specifically for nodes of some groups involved in rapid diversifications that historically have been problematic to resolve^2,29^. In particular, Maxillariinae+Stanhopeinae and Catasetinae+Cyrtopodiinae are now both strongly supported (LBP=100). In addition, our results also support the placement of *Dipodium* (Dipodiinae) is supported as sister to the rest of Cymbidieae, a relationship which was previously recovered from a few loci^25^. However, our plastid phylogenomic framework is still incomplete due to absence of representatives of Eriopsidiinae and Coleopsidinae.

One other novelty of our study is the inference of relationships in Zygopetalinae, a subtribe in which relationships have previously been poorly understood^56^. The most extensively sampled analysis of Zygopetalinae inferred from plastid markers (*mat*K-*ycf1*) ^29^ included 60 species and 27 genera, but relationships between most genera attained only low support. Our expanded molecular, but taxonomically reduced, matrix (i.e. 20 genera and 21 species) produced greater support for the backbone relationships in the subtribe, including the radiation of the *Huntleya* clade (*Dichaea, Huntleya, Chaubardia* and the *Chondrorhyncha* complex^56,57^). Nonetheless, relationships between the *Huntleya* grade (i.e. *Huntleya* clade + *Cryptarrhena*) and the remainder of Zygopetalinae still remains unresolved.

Our phylogenetic analyses further place for the first time in the orchid tree of life *Cheiradenia* and *Hoehneella* with moderate to strong support (Fig. 1, S1). *Cheiradenia* is a monospecific genus restricted to the lowland wet forests of Venezuela and Guyana whereas *Hoehneella* includes two species exclusively distributed in the Brazilian evergreen wet forests of the Brazilian states of Espirito Santo and São Paulo^58^. Referring to the similarity of both vegetative and floral reproductive characters, Pupulin ^58^ hypothesised that *Cheiradenia* should be closely related to members of the *Zygopetalum* clade (e.g. *Koellensteina, Paradisanthus*), with *Hoehneella* being related to the *Huntleya* clade (i.e. *Huntleya* and *Chaubardia*). Our ML tree supports both assumptions, placing *Cheiradenia* as sister to *Paradisanthus* with maximum support and *Hoehneella* as sister to *Chaubardia* in a moderately supported clade (83 LBP: Fig. 1, S1). *Koellensteina kellneriana* (the taxonomic type of the genus) clustered with *Acacallis* and not with *Otostylis* and *Paradisanthus*, and therefore we confirm that *Koellensteina* in the strict sense is related to *Acacallis*. In addition, *Otostylis* is recovered as sister to *Warrea* and not to *Paradisanthus* as previously suggested by Williams et al.^56^ based on a weakly supported placement. Our results also highlight the extensive and independent terrestrial and epiphytic habit transitions occurring in this clade, as most sister genera shows different habit types.

### 3.5. Novel and robust relationships in the most rapidly diversifying subtribe Pleurothallidinae

One of the most spectacular Neotropical plant diversifications is perhaps that of the Pleurothallidinae, for it involves the evolution of ∼5,000 species that have conquered virtually all biogeographical regions in the American tropics^32,45^. The rapid radiation of Pleurothallidinae occurring in the last ∼20 Myrs^28^ is associated with the evolution of a diverse suite of pollination systems ranging from food deception^59^ to pseudocopulation^60^ linked to dipterans^61,62^ and a complex array of reproductive and vegetative morphologies^22,32^. Understanding of relationships in the subtribe has relied mostly on relatively small number of markers^63–65^, which have informed with some confidence the phylogenetic placement and monophyly of genera in Pleurothallidinae, yet basal nodes in these trees have often lacked good support.

Several attempts have been conducted to estimate generic relationships in the subtribe, most of which have relied on nuclear rITS and plastid *matK* markers^66^. A synthesis of the phylogenetic relationships in the subtribe based on such studies was conducted by Karremans^67^. Here, a cladogram depicting the commonest topologies of relationships between genera was provided and nine clades were defined (termed “affinities” by the author) but without considering the magnitude of the support for these (see Figure 2 in Karremans^67^). Our plastid phylogenomic analysis recovered well-supported relationships in Pleurothallidinae that are mostly in line with previously published studies ^28,63,68^. However, these previous trees based on a handful of DNA nuclear and plastid markers yielded poor resolution and low support for backbone nodes as well as infrageneric relationships. In contrast, our plastid phylogenomic inferences recovered high support along the backbone, thus recovering novel placements. Some of these noteworthy well-supported relationships are the position of *Acianthera* as sister to *Myoxanthus* and *Dresslerella* as sister to *Barbosella*+*Restrepia* (Fig. 1, S1).

*Acianthera* includes over 300 species distributed throughout the American tropics and subtropics^64,69,70^, is often retrieved as sister to the remainder of Pleurothallidinae with moderate support^68^. Karremans^67^ used a series of “affinities” to describe to groups of genera affiliated with a core genus of these group and thus described the “*Acianthera* affinity” as the frequent clustering of several Central American genera with *Acianthera*^64^. Our study contradicts Karreman’s^67^ concept of the *Acianthera* affinity by placing with high support *Acianthera* in the *Restrepia* affinity as sister to *Myoxanthus. Dresslerella* was previously recovered with low support as sister to the remaining genera in the *Restrepia* affinity (*Barbosella, Echinosepala, Myoxanthus, Restrepia, Restrepiella* and *Restrepiopsis*). In contrast, our analysis robustly places *Dreslerella* as sister to *Restrepia* and *Barbosella*, a result that does not support the monophyly of the *Restrepia* affinity.

Although estimates of the ancestral distribution of the Pleurothallidinae are still uncertain, most of the early divergent Pleurothallidinae and their sister groups are found in the Antilles or Brazil^28^. The remarkable relationship recovered here for *Acianthera*+*Myoxanthus* could yield more clues about the biogeographic history and evolution of the subtribe because Brazil harbours a high species diversity of *Acianthera* and some of the early divergent clades in *Myoxanthus* (particularly the species close to *M. lonchophyllus*), whereas *Myoxanthus* is notably absent in the Antilles. In addition, other early divergent clades such as *Octomeria* and *Barbosella* are more diverse in Brazil. These early diverging clades share the lack of stem annulus as a morphological symplesiomorphy, a character that later appears in more diverse clades such as *Masdevallia+Dracula, Lepanthes*, and *Pleurothallis+Stelis*^71^. Members of these clades probably diversified after a migration to the mountainous areas of the northern Andes ca 16 ± 5 Ma and together account for almost 80% of the species in the subtribe^28^. The modern range extends mostly along the Andean and Central American mountain ranges. Here, another noteworthy relationship is that the less diverse *Specklinia* clade (*Scaphosepalum*+*Platystele*) was recovered as sister to the most species-rich clades of the subtribe (*Masdevallia, Lepanthes*, and *Pleurothallis*). In previous phylogenetic analyses *Specklinia* clade was recovered as sister to just *Pleurothallis*^28^.

Likewise, relationships between early divergent members in the *Lepanthes* affinity (*Anathallis, Draconanthes, Epibator, Lepanthes, Opilionanthe, Trichosalpinx* and *Tubella*) were largely weakly supported, demonstrating the need for increased taxon sampling, principally in *Lepanthopsis* and *Tubella*^32^. In particular, the early diversification of the *Lepanthes* affinity (>1500 spp.), inferred to have occurred around 8 Ma, has been linked to colonisation of newly formed environments in the Andean Cordillera, a product of accelerated mountain uplift and specific pollination systems (pseudocopulation and food mimicry^60^).

Another novel placement concerns *Teagueia* (diverse in Colombia, Ecuador and Peru^72–74^), which resembles *Platystele*^75^. Karremans^76^ had suggested a close relationship between *Teagueia* and *Scaphosepalum*, but our results place *Teagueia* as sister to *Platystele* with high support, thus corroborating the long-standing hypotheses of their sister relationship based on the of their reproductive structures^74,75^.

### 3.5. Evolutionary relationships in Orchidoideae

Our study provides a well-supported tree for Orchidoideae. Our ML inference supports the findings of Pridgeon et al.^35^ in which Diurideae is sister to Cranichideae and Codonorchideae to Orchideae. Our findings differ from Givnish et al.^2^ and Salazar et al.^34^, in which Diurideae/Cranichideae are sister to Codonorchideae, with Orchideae sister to all these (Fig. 2). Givnish et al.^2^ included all four tribes but only six of 21 subtribes of Orchidoideae, and the relationship of Diurideae to Cranichideae was poorly supported.

## Conclusions

This study presents a well-resolved, more densely sampled and strongly supported analysis of Orchidaceae and their absolute times of divergence than all previous such studies. For deep branches and recent diversifications in Cymbidieae and Epidendreae, support is improved, yet several recalcitrant nodes that historically have been challenging to resolve were also found (e.g. early divergent taxa in the Epidendroideae, initial radiation of the *Lepanthes* affinity in Pleurothallidinae). Similarly, our analyses provide the a well-supported result for Orchidoideae. Although taxon sampling was sufficient to resolve the relationships between the major clades in the family, sampling of unrepresented genera and representatives of Eriopsidiinae, Goodyerinae, and Coleopsidinae would further enhance our understanding of phylogenetic relationships.

## Material and methods

### 2.1 Sampling, DNA extraction and sequencing

Two-hundred and sixty-four species representing 117 genera, 28 subtribes and 18 tribes were sampled in this study. For 51 species plastid genomes were sequenced. Table S5 provides accession numbers of plastid genomes sourced from NCBI and GenBank numbers of those newly generated. Fresh leaves were stored in silica gel for subsequent DNA extraction using a CTAB method^77^. Total DNA was purified with silica columns and then eluted in Tris-EDTA^78^. DNA samples were adjusted to 50 ng/uL and sheared to fragments of approximately 500 bp.

#### High-throughput sequencing

The library preparation, barcoding and sequencing (Illumina HiSeqX) were conducted at Rapid Genomics LLC (Gainesville, FL, USA) and Genewiz GmbH (Leipzig, Germany). Pair-end reads of 150 bp were obtained for fragments with insert size of 300-600 bp. Overhangs were blunt ended using T4 DNA polymerase, Klenow fragment and T4 polynucleotide kinase. Subsequently, a base ‘A’ was added to the 3’ end of the phosphorylated blunt DNA fragments. DNA fragments were ligated to adapters, which have a thymine (T) overhang. Ligation products were gel-purified by electrophoresis to remove all unbound adapters or split adapters that were ligated together. Ligation products were then selectively enriched and amplified by PCR. For each sample, between one and 10 million paired-end reads were generated.

#### Plastid genome assembly and annotation

Raw sequences were quality filtered using Trimmomatic^79^ in order to eliminate sequencing artefacts, improve uniformity in the read length (>40 bp) and ensure quality (>20) for further analysis. Filtered sequences were processed with BBNorm^80^ to normalize coverage by down-sampling reads over high-depth areas of the genomes (maximum depth coverage 900x and minimum depth 6x). This step creates a flat coverage distribution in order to improve read assembly. Subsequently, overlapping reads were merged into single reads using BBmerge^81^ in order to accelerate the assembly process. Overlapping of paired reads was evaluated with Flash^82^ to reduce redundancy. Merged reads were used to carry out the whole genome de novo assembly with SPAdes (hash length 33,55,77)^83^.

To produce contiguous, linear plastid genome sequences we relied on a refence-based and *de-novo* approaches. The reference based approach was conducted on MIRA v. 4^84^, a software that maps read data against a consensus sequence of a reference assembly (simple mapping). MIRA has been useful for assembling complicated genomes with many repetitive sequences^85–87^. MIRA produces BAM files as output, which were subsequently used to generate consensus sequences in SAMTOOLS^88^. We sourced 11 reference plastomes from the NCBI repository that represent related species, namely: *Cattleya crispata, Goodyera fumata, Masdevallia picturata, M. coccinea, Oncidium sphacelatum* and *Sobralia callosa*. The *de-novo* assembly approach relied on GetOrganelle^89^, using the recommended default settings for assemblies of green-plant plastid genomes.

Newly sequenced and datamined plastid genomes were annotated through the Chlorobox portal of the Max Planck Institute^90^. Sequences were uploaded as fasta files, and running parameters were established as follow: BLAST protein search identity=65%, BLAST rRNA, tRNA, DNA search identity=85%, genetic code = bacterial/plant plastid, max intron length=3,000, options= allow overlaps. *Apostasia wallichii, Masdevallia picturata, Oncidium sphacelatum, Sobralia callosa* and *Goodyera fumata* were set as the ‘Server Reference’ and *Cattleya liliputana* was set as the ‘Custom Reference’ for CDS and tRNA, rRNA, primer, other DNA or RNA specifications.

#### Phylogenetic analysis

A set of 78 plastid genes was used to reconstruct phylogenetic relationships in Orchidaceae. These were aligned^91^ using MAFFT 7^92^ and subsequently concatenated (proportions of missing data per species is provided on Tab. S5). This step was performed at the supercomputing centre APOLO, EAFIT University, Medellín, Colombia. Phylogenetic reconstruction based on maximum likelihood (ML) was implemented in RAxML v. 8.0^93^, using 1,000 bootstrap replicates and the GTR+GAMMA model. Absolute age estimation analyses relied on fossil and secondary calibration points, strict and molecular clocks and a birth/death model implemented in BEAST v. 1.8^94^. The fossil constraint was added to the MRCA of *Dendrobium* following Xiang et al.^95^ using a normal distribution with mean value of 21.07 and a standard deviation (SD) of 3.0. Following Givnish et al.^2^, the two secondary calibration points were added to the root of the tree and MRCA of the Orchidaceae, using a normal distribution and mean values of 123.48 (SD=2.0) and 90 (SD=2.0). Because dating analyses conducted on dozens of gene alignments and hundreds of terminals are extremely computationally greedy, we estimated absolute ages on the five most phylogenetically informative genes (see below) and by constraining the tree topology to the ML tree derived from RAxML. For each clock model, we conducted two MCMC analyses with 250 million generations each with a sampling frequency of 10000 generations. The convergence of the strict and relaxed molecular clocks parameters was confirmed on the software TRACER v1.6. (http://tree.bio.ed.ac.uk/software/tracer/). Maximum clade credibility trees were summarised from the MCMC trees in the program TreeAnnotator v.1.8. of the software BEAST.

#### Phylogenetic informativeness profiles

To estimate the phylogenetic informativeness (PI) of plastid genes we calculated the per-site and net values for each assessed locus with the HyPhy substitution rates algorithm for DNA sequences^96^ using in the web application PhyDesign http://phydesign.townsend.yale.edu/). The input files were the consensus ML ultrametric tree converted with the function *chronos* of the R-package APE (http://ape-package.ird.fr/) using an smoothing rate of 1 and a relaxed clock model, and the partitioned concatenated gene alignments.

## Supporting information

Figure S1. Detailed maximum likelihood tree of the orchid family inferred from 78 plastid genes.

Figure S2. Chronogram of the orchid family as inferred from a strict molecular clock and a birth-death model.

Figure S3. Chronogram of the orchid family as inferred from a relaxed molecular clock and a birth-death model

Figure S4. Phylogenetic informativeness (PI) of 78 plastid gene alignments used in this study to infer orchid relationships

## Acknowledgments

We would like to thank Esteban Urrea for helping with the bioinformatics pipelines. We thank Norris Williams and the late Mark Whitten (University of Florida) for collecting and preparing the specimens. Kurt Neubig from Southern Illinois University provided the sequences of 11 new samples. We also thank Janice Valencia for critical feedback on the paper, Juan David Pineda Cardenas for advising about computational resources used through EAFIT and Juan Carlos Correa for computational advices at BIOS. The University of Costa Rica provided access to the genetic material for the projects B8257 and B6140. Finally, we would like to thank IDEA WILD for supporting with photographic equipment and Sociedad Colombiana de Orquideología for supporting M. A. Serna-Sánchez with a grant to conduct her undergraduate studies. O.A.P.E. is supported by the Swiss Orchid Foundation and the Lady Sainsbury Orchid Fellowship. A.A. acknowledges financial support from the Swedish Research Council (2019-05191), the Swedish Foundation for Strategic Research (FFL15-0196) and the Royal Botanic Gardens, Kew.

## Author Contributions Statement

M.A.S.S., O.A.P.E., and T.A. designed research; O.A.P.E., M.A.S.S., T.A., C.H. and A.A. generated new data; M.A.S.S., O.A.P.E., M.F.T., A.C.A.Y., J.A. and S.D. performed all analyses; O.A.P.E., D.B., M.W.C., M.A.S.S., S.D. and T.A. wrote the manuscript, with contributions from all authors.

## Legends

**Table 1**. Absolute ages and confidence intervals of main orchid lineages as inferred under a strict and relaxed molecular clocks and a Birth-Death model.

## Supplementary materials

**Table S1**. Detailed absolute ages, confidence intervals and rates for the orchid family as inferred under a strict molecular clock and a birth-death model. The table contains 528 rows and 16 columns and is available at https://doi.org/10.6084/m9.figshare.13185008.v1.

**Table S2**. Detailed absolute ages, confidence intervals and rates for the orchid family as inferred under a relaxed molecular clock and a birth-death model. The table contains 528 rows and 16 columns and is available at https://doi.org/10.6084/m9.figshare.13185008.v1.

**Table S3**. Phylogenetic informativeness per-site.

**Table S4**. Phylogenetic net informativeness.

**Table S5**. Voucher information and proportion of missing data in gene alignments.

**Figure S1**. Detailed maximum likelihood tree of the orchid family inferred from 78 plastid genes. LBP <100 are shown at nodes, with LBP <85 highlighted in red together with their corresponding subtending branches.

**Figure S2**. Chronogram of the orchid family as inferred from a strict molecular clock and a birth-death model. LBP at nodes <85 are highlighted in red together with their corresponding subtending branches. Blue bars at nodes denote 95% high density probability (HDP) absolute age intervals.

**Figure S3**. Chronogram of the orchid family as inferred from a relaxed molecular clock and a birth-death model. LBP at nodes <85 are highlighted in red together with their corresponding subtending branches. Blue bars at nodes denote 95% high density probability (HDP) absolute age intervals.

**Figure S4**. Phylogenetic informativeness (PI) of 78 plastid gene alignments used in this study to infer orchid relationships. A) Chronogram of Orchidaceae as inferred by PATH8 from the ML tree derived from RAxML; B) Per-site PI; C) Net PI.

