## Supplementary figures and images for "Plastid phylogenomics resolves ambiguous relationships within the orchid family and provides a solid timeframe for biogeography and macroevolution"

### Figure S1. Detailed maximum likelihood tree of the orchid family inferred from 78 plastid genes.

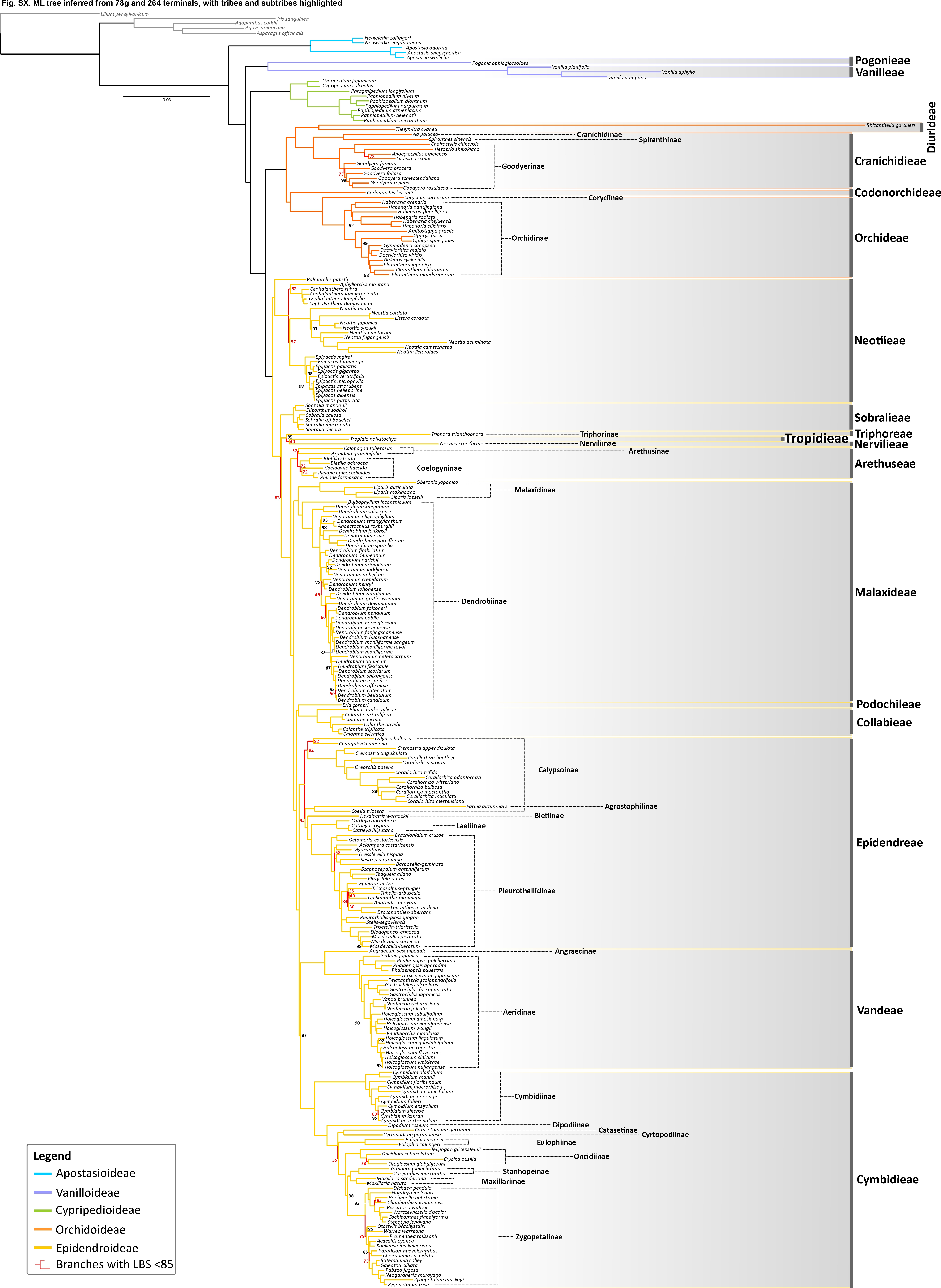

### Figure S2. Chronogram of the orchid family as inferred from a strict molecular clock and a birth-death model.

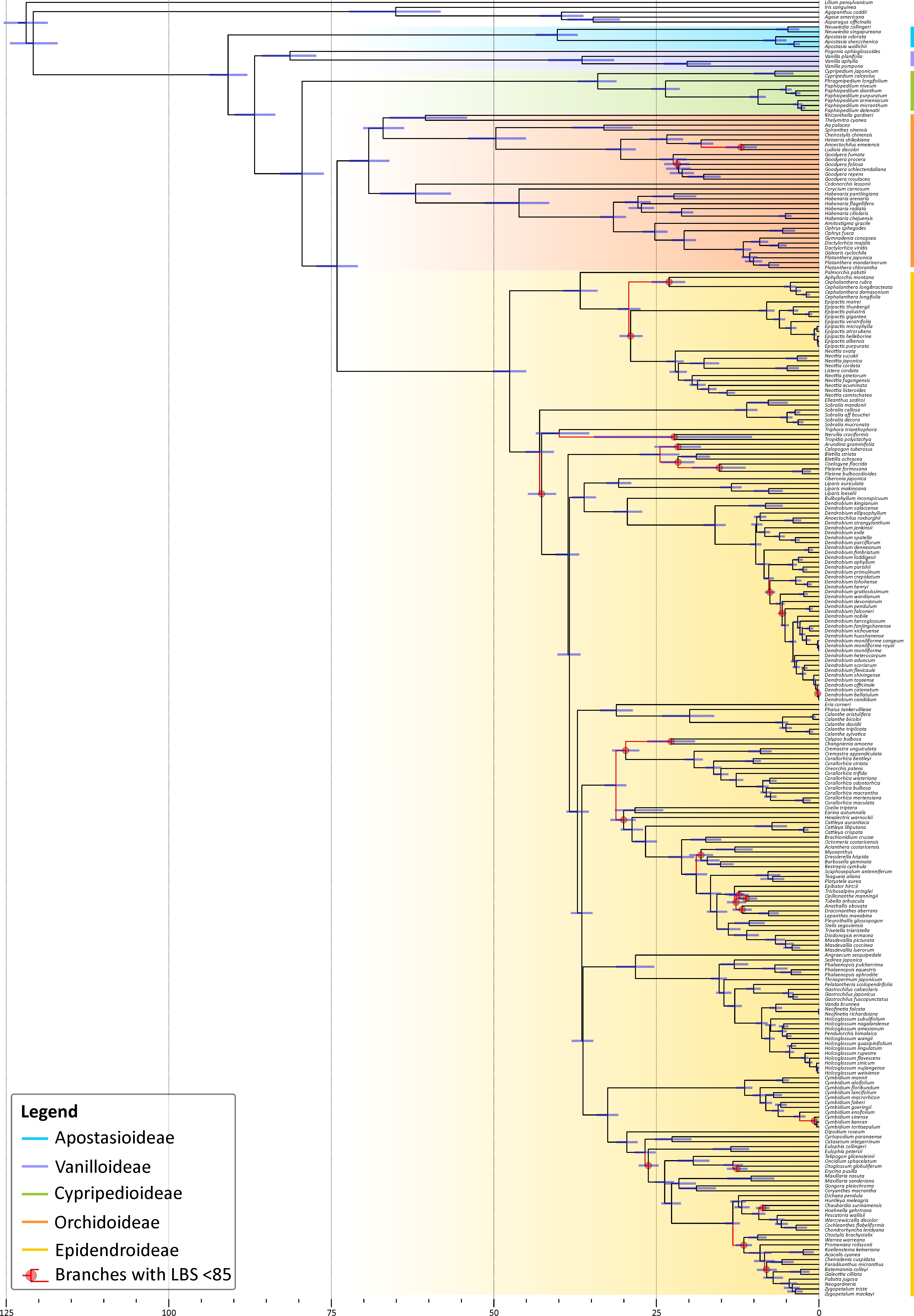

### Figure S3. Chronogram of the orchid family as inferred from a relaxed molecular clock and a birth-death model

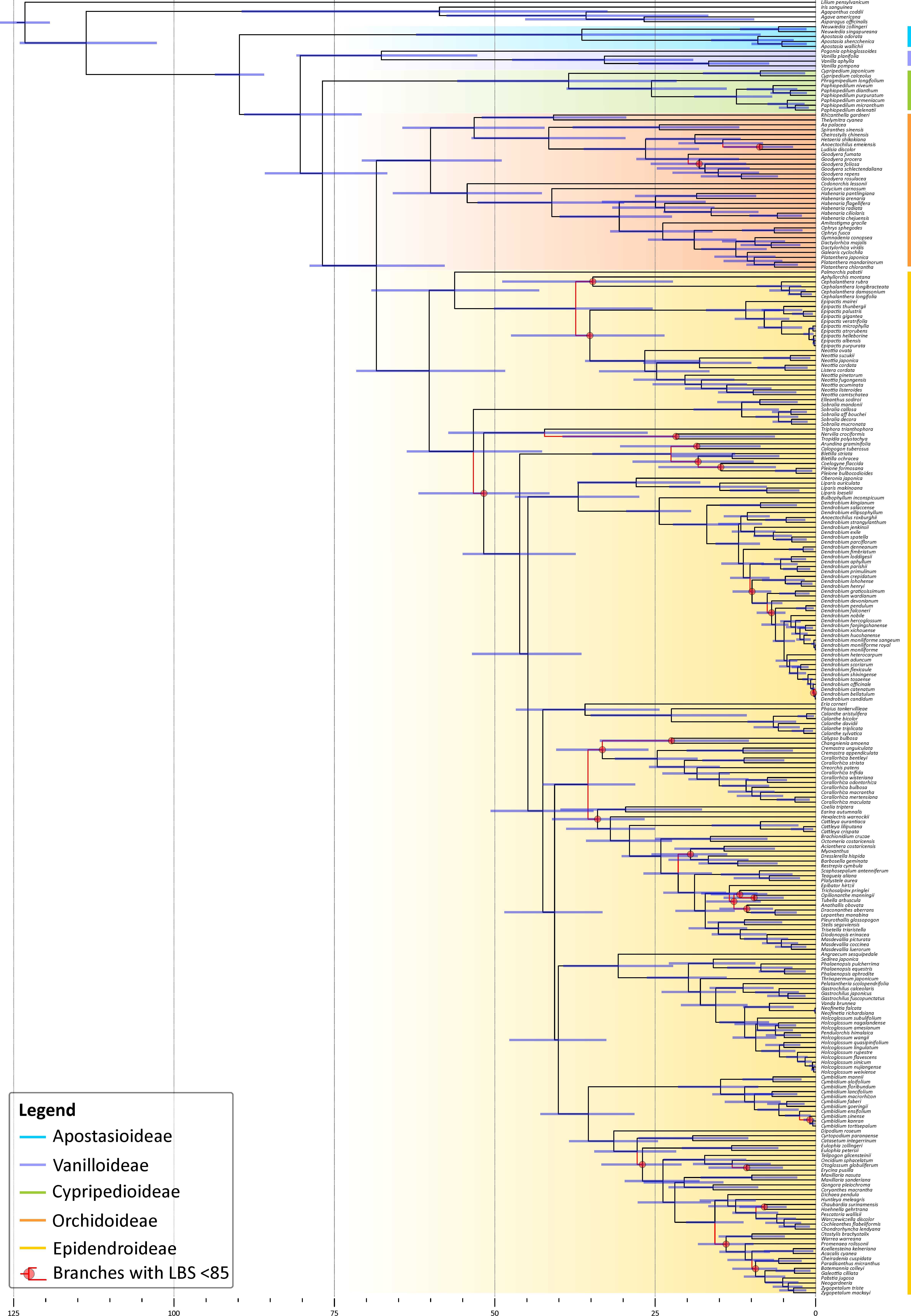

### Figure S4. Phylogenetic informativeness (PI) of 78 plastid gene alignments used in this study to infer orchid relationships

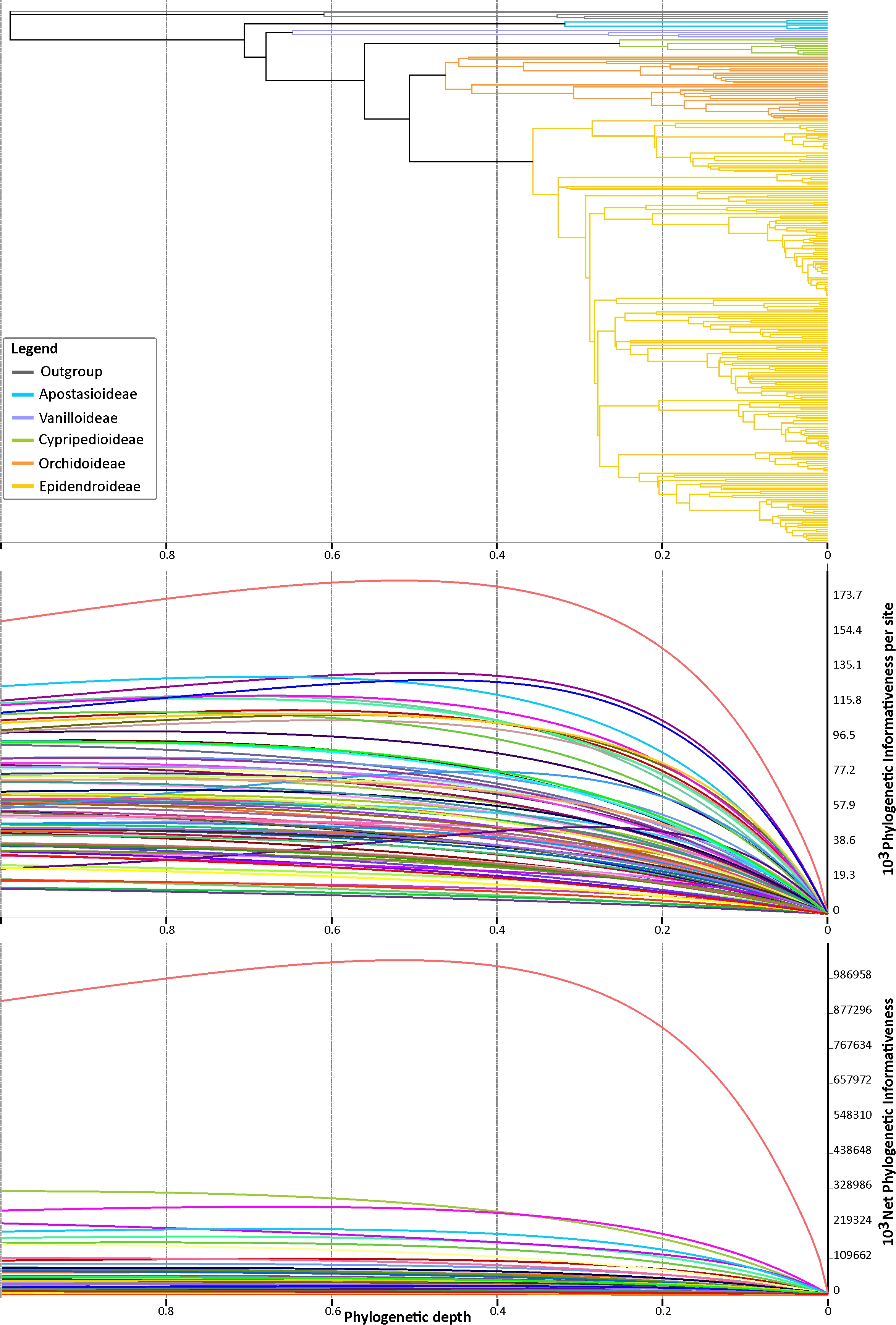
